# Ultrasound-based prediction of spawning time in females of Octopus vulgaris

**DOI:** 10.1101/2025.03.28.645890

**Authors:** L. Márquez, P. Weterings, S. Saber, B.C. Felipe, S. Maccuro, M. Serrano, E. Seuntjens, F. Acosta-Cifuentes, M. González, E. Almansa

## Abstract

There has been a great interest in common octopus *Octopus vulgaris* aquaculture over recent years and commercial production is coming within reach. In addition, *O. vulgaris* is a short-lived semelparous species with a long spawning period. Therefore, monitoring ovary maturation in cultured octopuses can be useful in the scheduling of hatchery operations and management of culture resources. The present work aimed to assess the ovary maturity status in cultured and wild common octopus by means of a noninvasive technique, ultrasonography, to anticipate the time of spawning in aquaculture. A total of 27 octopuses (15 cultured and 12 wild ones) in different ovarian maturation stages were used in ultrasonographic procedures. Ultrasound-based ovary volumes were compared with real ovarian volumes and then the ultrasonography-based gonadosomatic index (GSIU) was estimated. Data from cultured females indicate that GSIU was initially below 0.3 cm^3^ per 100 g of body weight in immature animals, and attained a fast increase until reaching about 4.0 cm^3^ per 100 g of body weight right before spawning. The steep increase in relative volume occurred similarly in females of different body weights. In addition, the specific growth rate of cultured animals decreases with time, and with GSIU. Lastly, a linear mixed model indicated a significantly negative and logarithmic relationship between the time to spawning and GSIU in captive females. That relationship will be useful to predict spawning time in immature and maturing females, helping in the management of octopus broodstocks in research and production facilities.

## Introduction

The common octopus *Octopus vulgaris* Cuvier, 1797, is an important commercial cephalopod, and it is presently under intensive research to develop its industrial production. The lifespan of this semelparous species is estimated to be between twelve and fifteen months (Katsanevakis & Verriopoulos, 2006). Juvenile specimens of the common octopus easily acclimate to farming conditions and can attain fast growth (Sánchez et al., 2014), and the survival and growth of the paralarval stage have been recently improved to values suitable for commercial production (Almansa et al., 2025). In addition, the manipulation of the duration of embryonic development, usually taking several weeks, helps researchers and potential farmers to manage resources for larval culture (Márquez et al. 2023). However, no similar knowledge is available to predict the spawning time of octopus females. The prediction of embryonic development duration, and the estimation of female spawning time would be particularly useful to schedule the use of tanks and other resources for the culture of octopus early stages. One potential way to produce suitable estimations of the spawning time is by monitoring of ovarian maturation stage. The gonad maturation in the genus *Octopus* is influenced by environmental factors like temperature, photoperiod and food availability (Forsythe and Hanlon, 1988; Mangold, 1983). At the same time, maturation is regulated by neuro-endocrine secretions that integrate the effects of external factors (Polese et al., 2015; Di Cristo, 2013; Wells and Wells, 1959). Regardless of the factors shaping the process, the stage of maturation in wild females can be assessed by calculating the gonadosomatic index (GSI), which is a measure of the relative volume of the gonad (Sieiro et al., 2020, 2014). As such, the GSI can be an indicator of the reproductive status also in cultured specimens of *O. vulgaris*, particularly if the procedure does not require animal slaughtering, as with biopsy or ultrasonographic techniques.

Considering the research on fish reproduction as a reference, ovarian biopsy (Du et al., 2017; Coward and Bromage, 2001), blood sex hormones (Mlingi et al., 2023; Du et al., 2017; Evans et al., 2004), and ultrasonographic imaging (Novelo and Tiersch, 2012) or, more recently, machine-learning-guided interpretation of ultrasonographic imagery (Yari et al., 2024), are all non-destructive methods chosen for assessing the maturation status. Among these non-destructive procedures, ultrasonography has the advantage of being a non-invasive technique. Ultrasound images of gonads of fish and gastropod females are suitable to measure primary biometric variables like ovary cross-section area, circumference and diameter (MacGarvey et al., 2021; Naeve et al., 2018; Jennings et al., 2005; Mattson, 1991), and ovary antero-posterior length (Naeve et al., 2018; Whiteman et al., 2005). Some authors provide derived biometric variables such as ovary volume, which requires a model for ovary shape (Dasgupta et al., 2024; Bureau du Colombier et al., 2015; Bryant et al., 2007) or machine-learning techniques (Yaris et al., 2024) and, subsequently, volumetric estimations of the GSI (Dasgupta et al., 2024). In general, ultrasonographic variables of fish ovaries show a good correlation with the same variables measured on the dissected organ (Naeve et al., 2023; Mattson, 1991).

In relation to non-destructive methods potentially useful to study the maturation process in cephalopods, several facts should be taken into account: i) there is a lack of publications about safe procedures to implement needle biopsy (Ponte et al., 2017), ii) the chemical nature of sex steroids is still under active research (Wang et al., 2022; Minakata and Tsuisui, 2016), and iii) ultrasound imagery with anesthesia is classified as a mild research procedure (Cooke et al., 2019), and has been used to study digestive tract, brain and arm tissue architecture of cephalopod mollusks (Berrueta et al., 2025; Ponte et al., 2017; Margheri et al., 2011; Grimaldi et al., 2007), as well as reproductive condition in gastropods mollusks (Boles et al., 2023; Zou et al., 2024).

The present work aims to characterize gonadal maturation in *O. vulgaris* females to anticipate the time of spawning with application to aquaculture and, also, to manage the breeding stock. Ultrasonography has been evaluated as a technique to monitor temporal changes in gonad size, and to estimate an ultrasonography-based gonadosomatic index (GSIU). These data allowed us to investigate the relationships between ultrasonographic gonadosomatic index (GSIU) and the time to spawning (t_Spawn_), as well as the specific growth rate (SGR) in cultured female specimens.

## Material and Methods

### Ethical approvals for cultured and wild animals

All animal experiments with cultured animals were performed in accordance with Spanish law 53/2013 in the framework of the European Union directive on animal welfare (Directive 2010/63/EU) for the protection of animals used for scientific purposes, following the Guidelines for the care and welfare of cephalopods in research proposed by (Fiorito et al., 2015). The described study was approved by the Ethics Committee of the National Competent Authority (Canary government, project number CEIBA 1377-2023 and CEIBA 1610-2024). Ethical approval was not required for protocols with wild animals, as all wild animals in this study originated from commercial fisheries.

### Sample collection, sampling area and maintenance in captivity

Cultured animals. A total of 15 females were caught by artisanal fishery (trap gear) in Tenerife waters (Canary Islands) from September 2023 to September 2024. Those individuals were maintained in 4,000 L rectangular fiber-glass tanks at the Tenerife Culture Plant (IEO-CSIC) following the conditions described in Reis et al. (2021). The tanks were maintained under a natural photoperiod for the geographical location (between 10L:14D to 14L:10D). Temperature and salinity were 22.6 ± 1.1°C and 36.8 ± 0.1 PSU respectively. Temperature was measured with a Tinytag Plus 2 thermometer (TGP-4020; Gemini Data Loggers Ltd., Tynitag West Sussex, United Kingdom) and salinity with a Refractometer S/Mill-E (ATAGO). About 50% of each tank surface was covered with a shady green net. The water flow of the rearing tank was 5 L/min. A mixture of frozen food based on squid (*Loligo gahi*) and crab (*Callinectes sapidus*) were given *ad libitum* to the broodstock each day. PVC pipes and plastic box were placed inside the tanks to provide 2–3 dens for each specimen. The presence of eggs was verified on a weekly basis to avoid disturbing the breeders.

Wild animals. A total of 51 wild females were obtained from landings from artisanal (pots) and trawl fisheries in the Mediterranean Andalusian waters, between March and May 2024. Fresh octopuses were analyzed in a laboratory where the following measurements were recorded: total weight (TW) (to the nearest 0.1 g), dorsal mantle length (to the nearest mm), ovary weight (to the nearest 0.01g), and ovary volume (to the nearest cm^3^) measured by the water-displacement method according to Scherle (1970) and Saber et al. (2015). A total of 12 out of the 51 females were subjected to ultrasonographic procedures to obtain ultrasound-based estimations of ovary volume (see below).

### Anaesthesia procedure for cultured common octopus

The common octopus was gently removed from its home tank using a net, subsequently transferred to a vessel with a secure lid, as they often try to escape (Andrews et al., 2013), and submerged in aerated seawater containing 1.5 – 2% absolute ethanol (EtOH) (PanReac, AppliChem) which is an effective anaesthetic for *O. vulgaris* according to Fiorito et al. (2015). During each sampling procedure, the anaesthetized individual was held and the ovary was scanned and the TW was recorded (to the nearest 0.1 g). The octopuses were then immediately returned to water, where their recovery was monitored.

### Ultrasonographic procedures, ovary volume, gonadosomatic index, and specific growth rate in cultured female octopuses

To obtain a scan, the octopus was positioned on its ventral side and an 8 Mhz linear echo sounder (Esaote Pie Medical model, Falco 100) was used. Ultrasonography-based ovarian volume was estimated using the biometric measurements obtained from ultrasound images (anteroposterior, dorsoventral and left-right axes), and the ultrasonography-based gonadosomatic index (GSIU) was subsequently calculated considering the ovary as an ellipsoid (Vol_U_ = (4/3) π r_1_ r_2_ r_3_, where r_1_, r_2_ and r_3_ are the three semi-axes). Female total body weight (TW) was also measured to the nearest 0.1 g. A volumetric, ultrasound-based gonadosomatic index (GSIU) was calculated for cultured females of common octopus as: GSIU (cm^3^ per 100 g) = 100 × Vol_U_ (cm^3^) / TW (g). In addition, the specific growth rate (SGR) was calculated as the percentage of TW increase per day, according to Reis et al. (2021): SGR = (lnTW_2_ – lnTW_1_) / time, where TW_1_ and TW_2_ are initial and final total weights for the period under consideration.

### Ultrasonographic procedures, ovary volume and gonadosomatic index in wild female octopuses

Ultrasonographic procedures in wild specimens from the Mediterranean coast were carried out with an ultrasound equipment (Mindray M9 VET) and a L12-4S linear array to scan ovaries in the same position as in the case of cultured octopuses. The frequency range was 6.6-12 Mhz.

### Statistical analyses

The relationship between the ultrasonography-based ovarian volume and real ovarian volume in wild females were analyzed by means of least squares linear regression. The point corresponding to the female with the largest ovarian weight and, consequently, the largest volume, was below the general regression line for the other 11 females. Since the ultrasonographic estimation of volumes of large organs can be limited by the size and type of the ultrasound transducer (MacGarvey et al., 2021), this ovary was subsequently excluded from the regression.

The relationship between GSIU and time to spawning in cultured females was analyzed by means of a linear mixed model (LMM), with individual as the cluster variable. Time to spawning was selected as a dependent variable and GSIU as a covariate. Since the visual inspection of the plot indicate curvilinearity, a second LMM with time to spawning as dependent variable and ln(GSIU) as covariate was run. For each LMM, intercept, slope, or intercept and slope were tested as random effects. The selected model was that one resulting in a non-singular solution and a lower AIC (Akaike Information Coefficient). Normality and homoscedasticity of residuals were checked in a Q-Q plot, and a residuals-predicted values plot, respectively. Given that it was possible to measure GSIU and time to spawning in 10 females, that sub-set of points was analysed by means of the LMM.

To study the relationship between the specific growth rate (SGR) and GSIU, SGR values were plotted against the mean GSIU obtained in the same period used to calculate SGR. When a LMM was fitted, only singular solutions were obtained, thus, this type of analysis was ruled out. As an alternative, the ratio between the increment in SGR (between the last and the first measurement), and the increment in GSIU (also between the last and the first measurement) was calculated for each individual. This ratio was calculated in individuals with two or more SGR estimations (i.e., eight individuals). The ratio was subjected to a one-sample t-test against a zero value. A mean ratio below 0 would indicate that SGR decreases with GSIU, a mean ratio above 0 would indicate that SGR increases with GSIU, at last, a mean ratio not significantly different from 0 would indicate no relationship between SGR and GSIU. For statistical significance, significance level of 0.05 was assumed in all tests.

## Results

### Relationship between the ultrasonography-based ovarian volume and real ovarian volume in wild female common octopus

To benchmark ultrasonography as a valuable method to measure gonadal volume, we first compared ultrasonography measurements with actual volumetric measurements of dissected gonads obtained from Octopus vulgaris females from the wild. Two estimations of ovary volume were obtained: one estimation based on ultrasound images, and the other one based on the water-displacement method of Scherle (1970). Both estimations of ovarian volume were compared by regression techniques.

Figure 1 shows the relationship between the ultrasonographic-based ovarian volume (Vol_U_) (cm^3^) and the Sherle-based ovarian volume (Vol_Scherle_) (cm^3^) in 12 wild females of *Octopus vulgaris*. When Vol_Scherle_ was below 120 cm^3^, a linear correlation was obtained (n=11, R > 0.999, p < 0.001), so that Vol_U_ = 0.7408 Vol_Scherle_. As previously mentioned, the point with a Vol_Scherle_ close to 180 cm^3^ clearly stands below the regression line, and was excluded (Fig. 1).

**Figure 1.**
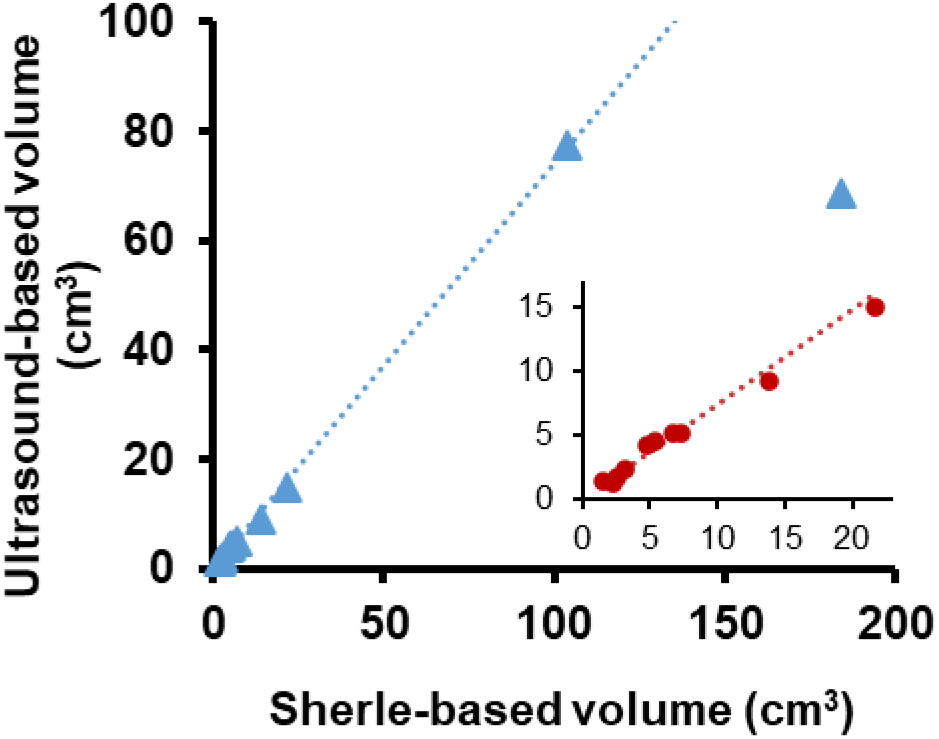
Relationship between the ultrasound-based volume and the Scherle-based volume of ovary in wild *Octopus vulgaris*. The inserted plot magnifies the regression line for Scherle-based volumes between 0 and 25 cm^3^.

### Relationship between ultrasonography-based gonadosomatic index (GSIU) and body wet weight (WW) in cultured females of Octopus vulgaris

Having established ultrasonography as a reliable method to measure gonadal size in a non-invasive manner, we next applied the method over a longer period of female maturation. The ultrasound images were used to monitor the temporal changes in ovary size of 15 cultured females. Ultrasonographic images of the ovaries were obtained every 15 to 30 days from capture to the time of spawning (period varying from a minimum of 26 days to a maximum of 101 days). Our study combined animals of different sizes at the onset of measurements.

Ten out of 15 females underwent a fast increase in GSIU, starting when total weight (TW) was approximately between 2 and 3 kg. Three animals showed a similar rapid increase in GSIU when TW was larger than 4 kg. At last, the rapid change in GSIU started at a weight below 2 kg in two females (Fig. 2). The starting value of GSIU was typically below 0.3 cm^3^ (100 g)^−1^. The last measured GSIU showed more variability, depending on the degree of gonad maturation, but the highest values were close to 4.0 cm^3^ (100 g)^−1^. These results show variability in the weight at which ovarian maturation begins, suggesting that maturation might be regulated independently from body weight.

**Figure 2.**
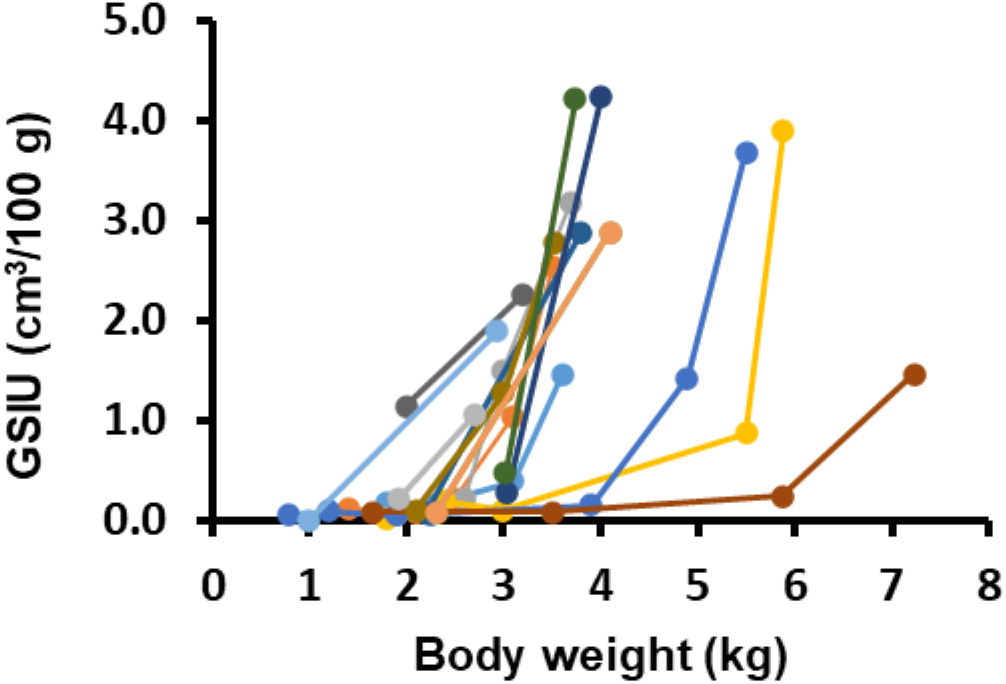
Changes in ultrasonographic GSIU with total weight in females of *Octopus vulgaris* reared in captivity. Different colors represent different individuals.

### Relationship between ultrasonography-based gonadosomatic index (GSIU) and time to spawning (t_Spawn_) in cultured females of Octopus vulgaris

As the end of maturation results in a spawning event, we hypothesized that GSIU measurements could provide a means to predict the spawning time. Plotting the GSIU values in relation to time to spawning, we observed that GSIU values increased from 0.1-0.2 cm^3^ (100 g)^−1^ for immature females to 4.0 cm^3^ (100 g)^−1^ in females on the verge of spawning. A negative correlation (r=-0.85, p<0.001) was observed between GSIU and time to egg laying (Fig. 3). The whole set of points suggests a curvilinear and concave relationship.

**Figure 3.**
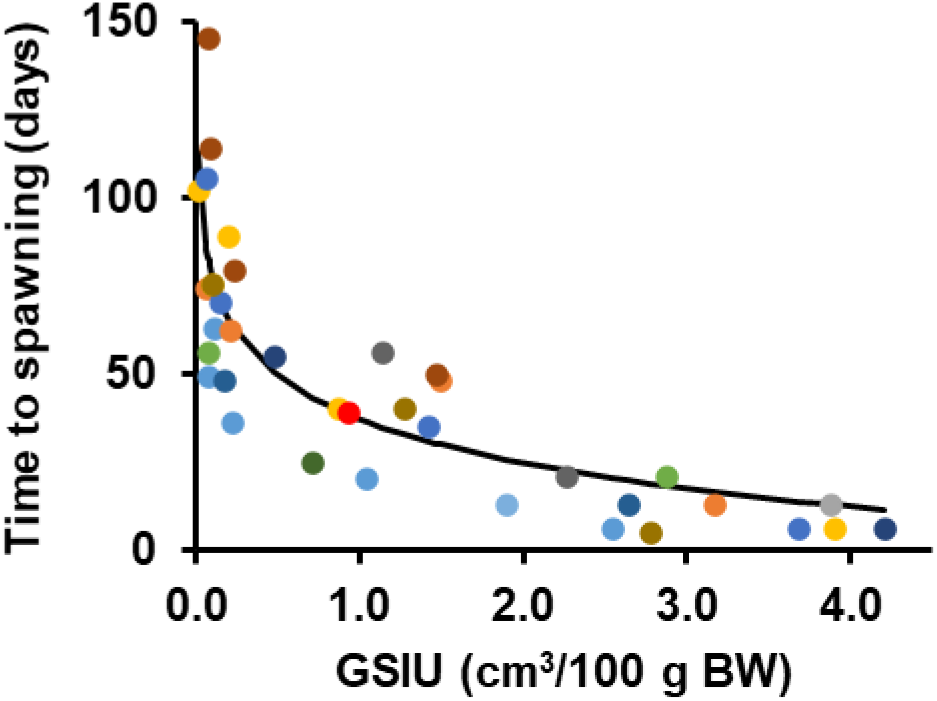
Relationship between gonadosomatic index (GSIU) and the number of days until the spawning event for each octopus. Different colors represent different individuals. The black line represents the regression line derived from a linear mixed model with a random intercept among individuals and fixed slope.

The comparison of Akaike Information Coefficient (AIC) for linear mixed models with different random effects (slope, intercept, or slope plus intercept) indicated that a model with random intercept and fixed slope was the most suitable. The fixed effect (GSIU) was highly significant (p < 0.001), and it explained about 68 % of the variance of the dependent variable (R^2^ Marginal = 0.684), whereas the combination of fixed plus random effects explained more than 83 % of the variance (R^2^ Conditional = 0.835). The mixed model resulted in the following equation to predict spawning: Time to spawning (days) = –17.8 ln(GSIU, cm^3^ (100 g)^−1^) + 37.2. As shown in Fig. 3, the relationship was curvilinear and concave, with the slope being more negative for GSIU values between 0 and 1. However, it became nearly linear for GSIU above 1.0. Animals with a GSIU value larger than 0.5 spawned within ~60 days.

### Relationship between specific growth rate (SGR) and ultrasonographic index (GSIU) of cultured females of Octopus vulgaris

Next, we wanted to use our data to calculate to what extent gonadal maturation was correlated or not with body growth. With this objective, the specific growth rate (SGR) between two consecutive ultrasonography sessions was calculated to monitor the specimen’s growth rate up to the final session before spawning. The relationship between GSIU and SGR is shown in Fig. 4. It shows a decrease in SGR with ovary size. The linear mixed model equation was not fitted in this case due to the production of singular solutions. However, the ratio ΔSGR / ΔGSIU was negative for all the individuals with more than one SGR estimation. The mean value of that ratio was −0.96 (Fig. 4b), and significantly different from zero in a t-Student test (n=8, t = −4.49, p = 0.003). Thus, a negative relationship between SGR and GSIU was apparent, indicating that growth slowed down when animals approached spawning.

**Figure 4.**
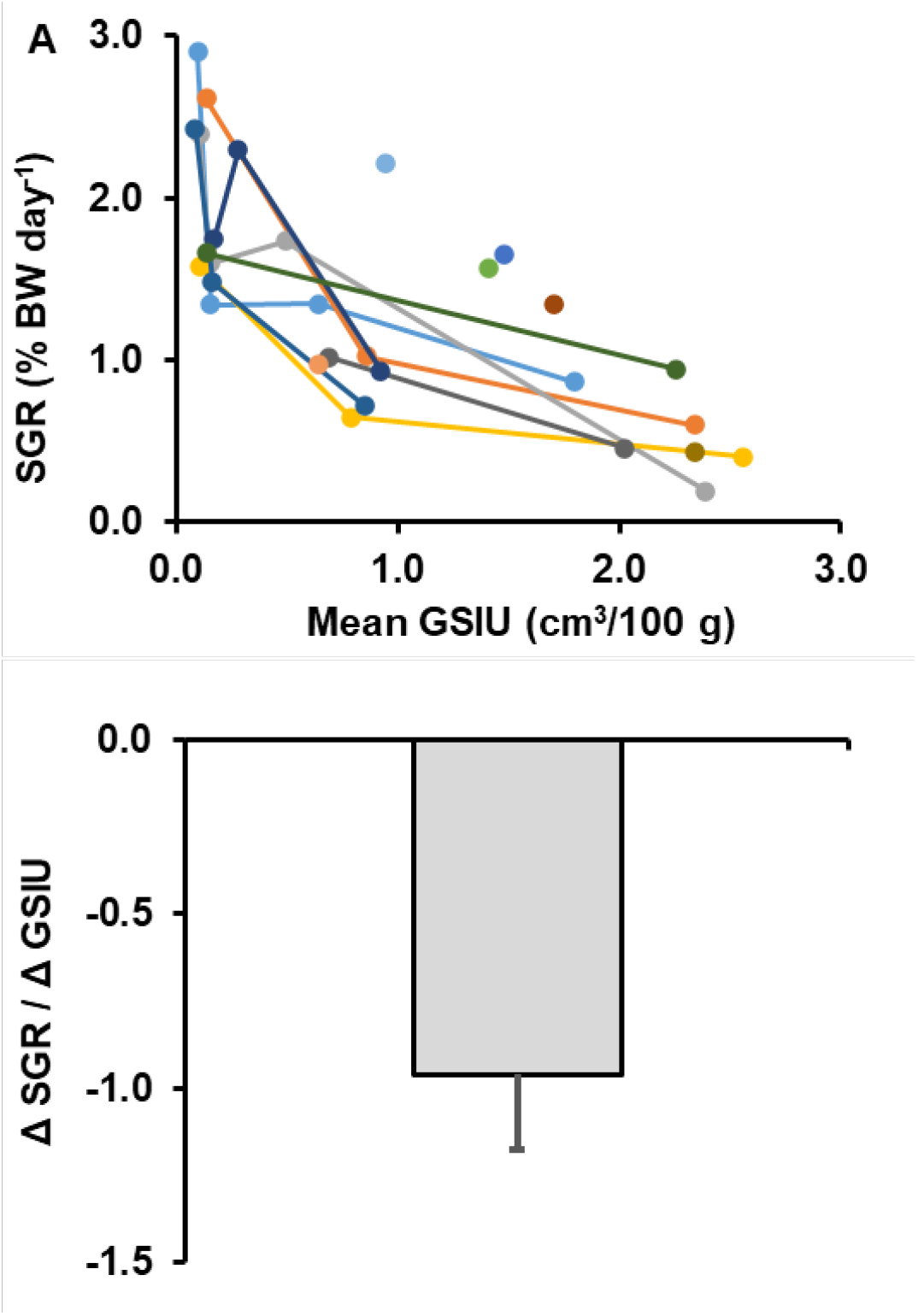
Relationship between GSIU (cm^3^/100 g) and SGR (% TW/ day). A. Plot of SGR versus mean GSIU for each individual. B. Mean and standard error of the ratio SGR/GSIU. Different colors represent different individuals.

## Discussion

The ultrasonography-based ovarian volume showed a good and positive correlation with the real ovarian volume of wild females. However, the regression line underestimated the real ovarian volume by approximately 25 %. Similarly, ultrasonography has also been reported to underestimate ovarian volume in fishes while showing a good correlation between estimated and real volumes (Dasgupta et al., 2024; Bryant et al., 2007). Inaccuracies of the shape model selected for the calculation of ovarian volume can produce deviations from real values. Researchers studying fish gonads frequently assume fusiform-like models for the shape of fish ovaries, since they are typically much longer than wider (Dasgupta et al., 2024; Bureau du Colombier et al., 2015; Macbeth et al., 2011; Bryant et al., 2007). In the case of common octopus, it is more reasonable to choose an ellipsoidal geometry as a model for ovary shape. However, when the ultrasound transducer is placed on the mantle, the ovary likely becomes flattened at the point of pressure, which implies that ovary shape would not exactly fit the ellipsoidal model during the ultrasonographic procedure. Ovary flattening due to transducer operation has already been reported in fish (McGarvey et al., 2021). In any event, it is clear that this effect does not necessarily affect the adequacy of the regression line, but only the value of the regression slope, thus it is not a serious problem since such a bias can be corrected.

Size estimation of large gonads by ultrasonography have been reported to result in underestimations depending on transducer type and size (Jennings, et al., 2005; McGarvey et al., 2021). This effect can be a more relevant problem because the bias would only affect large gonads in relation to the transducer, potentially affecting the graphic pattern displayed by data sets. In any case, this bias is limited to the more upper range of gonad sizes. In the present work, one ovary with an estimated volume above 180 cm^3^ stood out of the general regression line between ultrasonographic volume and real volume. More ultrasonographic measurements of octopus large ovaries (volume estimations clearly above 100 cm^3^) will be useful to ascertain if this type of gonads is systematically below the regression line for smaller ovaries. Nevertheless, the data presented herein highlight the utility of ultrasonographic estimates for ovaries at least up to 100 cm^3^ with the linear transducer L12-4S (used in wild octopuses), and supports a simple equivalence between estimated and real ovarian volumes if an ellipsoidal shape model is assumed (Fig. 1): ultrasonographic volume = 0.74 (real volume). Taking into account this volume equivalence, together with the mean ovary density found in wild females (n=47, 1.108 ± 0.041), the weight-based GSI can be estimated as approximately 1.5 times (1.108/0.74) the ultrasonographic volume-based GSI (GSIU).

The most important finding of the present work is a predictable and quantifiable negative relationship between the time to spawning (the period between ultrasonography time and spawning time), and GSIU. The physiological basis for this relationship rests on the increase in the number of oocytes, and the increase in oocyte size during ovary maturation (Cuccu et al., 2013a). The linear mixed model herein presented indicates that time to spawning decreased from approximately 40 to 10 days when the GSIU progressed from 1.0 to 4.0 cm^3^ (100 g BW)^−1^ or, after applying the 1.5 equivalence, when the real GSI (GSI = 100 × ovary weight (g) / total weight (g)) increased from 1.5 to 6.0 g (100 g TW)^−1^. The initial bended shape of the curve, as well as the large variation in time measurements when GSIU is small, roots in grouping pre-maturation females that have small gonads for prolonged periods, with those that are starting their maturation. When gonads are small (pre-maturation), prediction becomes unreliable. Further research should be focused on the regression line for females with GSIU above 2.0 (real GSI approx. above 3.0), when females have embarked on the maturation process, since the relationship seems to be linear of quasi-linear above that value. Further refinement of the regression can make measurement even more reliable, and in theory should make prediction of spawning time feasible after a single measurement in this quasi-linear period.

A remarkable finding was that maturation seemed to occur in a stereotyped manner, independent of the body weight of the female. Nearly all the females in the present work attained a fast increase in GSI at some point after being stocked in rearing tanks. This pattern with a fast change in GSI has been clearly obtained in wild females from the Tyrrhenian Sea during spring and summer, when the mean WW was above 1700 g, and after a peak of sex steroids in the ovary (Di Cosmo et al., 2001). Data on European populations of the common octopus support that weight at which 50 % of females reach maturity (WT_50_) ranges from 500 g to 1400 g in Mediterranean populations (Cuccu et al., 2013b; González et al., 2011; Tirado et al., 2003), and from 1100 to 2500 g in Atlantic populations (Sieiro et al., 2014; Lourenço et al., 2012; Otero et al., 2007; Carvalho and Reis, 2003; Tirado et al., 2003; Silva et al., 2002). Additionally, in Atlantic populations of Spain (Andalusian and Galician coasts) and Portugal, the percentage of mature females increases steadily in the weight interval 1500 – 3000 g, and 2000 – 3000 g, respectively (Sieiro et al., 2013; Lourenço et al., 2012; Silva et al., 2002). These data are consistent with our results from Canary Islands animals reared in captivity (Fig. 2), since the majority of females showed a steep increase in GSI in the range of weight 2000 – 3000 g, although a smaller percentage of octopuses underwent a similar rapid GSI change at body weights below 2000 g, or above 4000 g. Thus, we cannot rule out that a minimal body weight is necessary before gonadal and oocyte maturation can start, but once initiated, it follows a predictable course.

Present data support a statistically significant and negative relationship between the mean GSI of a female in a given period, and its SGR in the same time interval, in spite of the fact that inter-individual variability is also apparent. O’Dor and Wells (1978) reported reduced protein synthesis in arm muscles, and fast growth of ovary in females entering precocious maturity after removing brain sub-pedunculate lobes. This process could affect the specific growth rate in terms of body weight. Much more recent works (González et al., 2011, Otero et al., 2007; Rosa et al., 2004) sampled wild populations of *O. vulgaris* and looked for correlations between the condition of ovary, digestive gland or body muscles, as evidence of a trade-off between somatic and reproductive growth, but no such correlations were found. On the other hand, some authors (Sieiro et al., 2020) do not completely rule out a partial reallocation of energy from somatic to reproductive growth. In any case, with or without energy reallocation, it is not clear how ovary growth can depress the SGR in terms of whole body which includes the gonad. Further studies in this area could elucidate the origin of the nutrients that comprise the yolk, which might contribute to the enhancement of spawning quality and other related aspects.

In conclusion, these data show that the relationship between GSIU and spawning time can be applied to predict the start of egg laying in order to schedule future tasks and manage resources in experimental and production hatcheries. The relationship between time to spawning and GSIU can be particularly useful for octopus farmers since, in contrast to fish, octopus females can store spermatophores in the oviduct glands until they mature and gonad maturation don’t show a clear correlation with the female weight (Wells, 1978).

## Acknowledgements

This study has received funding from the European Maritime, Fisheries and Aquaculture Fund; ECOPHYN Project (PID2021-126824NB-C31-C33) funded by Spanish Ministerio de Ciencia e Innovación MICIU/AEI/10.13039/501100011033 and “ERDF/EU” and is part of the ThinkInAzul programme supported by MICIU, European Union NextGeneration EU (PRTR-C17.I1) and Gobierno de Canarias (SD2218/6897).

## References

Almansa, E., Márquez, L., Rosas, C., Martín, MV., Navarro, JC., Uriarte, I., Gestal, C., Fernández-Álvarez, FA., Gallardo, P., Varó, I., Farías, A., Cardenete, G., Caamal-Monsreal, C., Rodríguez-Barreto, D. and Morales, AE. In press. Octopods aquaculture: Reproduction, rearing technology, nutritional physiology, welfare and health status. In “Aquaculture and Living Resource Management – Volume 2”, edited by Prof. Leonel Pereira to be published by CRC Press (Taylor & Francis Group). ISBN 978-1-032-34632-8. In press.

Berrueta, M., López, A. V., Radonic, M., Goroso, B. G., Aristizabal, E., 2025. In vivo determination of sex and reproductive status of *Octopus tehuelchus* (Cephalopoda: Octopodidae) by ultrasound techniques. Marine and Fishery Sciences (MAFIS), 38(3).

Boles, S.E., Rofers-Bennett, L., Bragg, W.K., Bredvik-Curran, J., Graham, S., Gross, J.A., 2023. Determination of gonad reproductive state using non-lethal ultrasonography in endangered black (*Haliotis cracherodii*) and white abalone (*H. sorenseni*). Front. Mar. Sci. 10, 1134844.

Bryant, J.L., Wildhaber, M.L., Papoulias, D.M., DeLonay, D.J., Tillitt, D.E., Annis, M.L., 2007. Estimation of gonad volume, fecundity, and reproductive stage of shovelnose sturgeon using sonography and endoscopy with application to the endangered pallid sturgeon. J. Appl. Ichthyol. 23, 411–419.

Bureau du Colombier, S., Jacobs, L., Gesset, C., Elie, P., Lambert, P., 2024. Ultrasonography as a non-invasive tool for sex determination and maturation monitoring in silver eels. Fish. Res. 164, 50–58.

Carvalho, J.M.N., Reis, C.S., 2009. Contributions to knowledge on the maturation and fertility of the common octopus *Octopus vulgaris* Cuvier, 1797 on the Portuguese coast. Bol. Inst. Esp. Oceanogr. 19, 473–481.

Cooke, G.M., Anderson, D.B., Begout, M.L., Dennison, N., Osorio, D., Tonkins, B., Kristiansen, T., Fiorito, G., Galligioni, V., Ponte, G., Andrews, P.L.R., 2019. Prospective severity classification of scientific procedures in cephalopods: report of a COST FA1301 working group survey. Lab. Anim. 53, 541–563.

Coward, K., Bromage, N.R., 2001. Stereological validation of ovarian biopsy as a means of investigating ovarian condition in broodstock tilapia *in vivo*. Aquaculture 195, 183–188.

Cuccu, D., Mereu, M., Porcu, C., Follesa, M.C., Cau, A.L., Cau, A., 2013a. Development of sexual organs and fecundity in *Octopus vulgaris* Cuvier, 1797 from the Sardinian waters (Mediterranean Sea). Mediterr. Mar. Sci. 14, 270–277.

Cuccu, D., Mereu, M., Cau, A., Pesci, P., Cau, A., 2013b. Reproductive development versus estimated age and size in a wild Mediterranean population of *Octopus vulgaris* (Cephalopoda: Octopodidae). J. Mar. Biol. Assoc. U. K. 93, 843–849.

Dasgupta, S., Biswas, G., Tripathi, G., Pathan, A.G., Ghoshal, T.K., Singh, P.K., Jana, T., Mitra, P., Samanta, S., De, D., Adhikari, S., Manna, R.K., Sahoo, A.K., 2024. Assessment of sex, gonad volume, and reproductive maturation status of the Indian shad, *Tenualosa ilisha*, using ultrasonic imaging: a rapid and non-invasive tool. J. Appl. Ichthyol. 2024, 6597804.

Di Cosmo, A., Di Cristo, C., Paolucci, M., 2001. Sex steroid hormone fluctuations and morphological changes of the reproductive system of the female of *Octopus vulgaris* throughout the annual cycle. J. Exp. Zool. 289, 33–47.

Di Cristo, 2013. Nervous control of reproduction in *Octopus vulgaris*: a new model. Invert. Neurosci. 12, 27–34.

Du, H., Zhang, X., Leng, X., Zhang, S., Luo, J., Liu, Z., Qiao, X., Kynard, B., Wei, Q., 2017. Gender and gonadal maturity stage identification of captive Chinese sturgeon, Acipenser sinensis, using ultrasound imagery and sex steroids. Gen. Comp. Endocrinol. 245, 36–43.

Evans, A.F., Fitzpatrick, M.S., Siddens, L.K., 2004. Use of ultrasound imaging and steroid concentrations to Identify maturational status in adult steelhead. N. Am. J. Fish. Manag. 24, 967–978.

Forsythe, J.W., Hanlon, R.T., 1988. Effect of temperature on laboratory growth, reproduction and life span of *Octopus bimaculoides*. Mar. Biol. 98, 369–379.

González, M., Barcala, E., Pérez-Gil, J.L., Carrasco, M.N., García-Martínez, M.C., 2011. Fisheries and reproductive biology of Octopus vulgaris (Mollusca: Cephalopoda) in the Gulf of Alicante (Northwestern Mediterranean). Medit. Mar. Sci. 12, 369–389.

Grimaldi, A.M., Agnisola, G., Fiorito, G., 2007. Using ultrasound to estimate brain size in the cephalopod *Octopus vulgaris* Cuvier *in vivo*. Brain Res. 1183, 66.73.

Jennings, C.A., Will, T.A., Reinert, T.R., 2005. Efficacy of a high- and low-frequency ultrasonic probe for measuring ovary volume and estimating fecundity of striped bass *Morone saxatilis* in the Savannah River Estuary. Fish. Res. 76, 445–453.

Katsanevakis, S., Verriopoulos, G., 2006. Seasonal population dynamics of *Octopus vulgaris* in the eastern Mediterranean. ICES J. Mar. Sci. 63, 151–160.

Lourenço, S., Moreno, A., Narciso, L., González, A.F., Pereira, J., 2012. Seasonal trends of the reproductive cycle of *Octopus vulgaris* in two environmentally distinct coastal areas. Fish. Res. 127-128, 116–124.

Macbeth, B.J., Frimer, H.D., Muscatello, J.R., Janz, D.M., 2011. Use of portable ultrasonography to determine ovary size and fecundity non-lethally in northern pike (*Esox lucius*) and white sucker (*Catostomus commersoni*). Water Qual. Res. J. Can. 46, 43–51.

McGarvey, L.M., Ilgen, J.E., Guy. C.S., McLellan, J.G., Webb, M.A.H., 2021. Gonad size measured by ultrasound to assign stage of maturity in burbot. J. Fish Wildl. Manag. 12, 241–249.

Mangold, K., 1983. Food, feeding and growth in cephalopods. Mem. Natl. Mus. Vic. 44, 81–93.

Margheri, L., Ponte, G., Mazzolai, B., Laschi, C., Fiorito, G., 2011. Non-invasive study of *Octopus vulgaris* arm morphology using ultrasound. J. Exp. Biol. 214, 3727–3731.

Márquez, L., Martín, M.V., Larson, M., Almansa, E., 2023. Application of the thermal time model of embryonic development duration to the culture of octopod cephalopods. Aquac. Rep. 30, 101563.

Mattson, N.S., 1991. A new technique to determine sex and gonad size in live fishes by using ultrasonography. J. Fish Biol. 39, 673–677.

Mlingi, F.T., Puvanendran, V., Burgerhout, E., Tveiten, H., Tomkievicz, J., Kjørsvik, E., Mommens, M., 2023. Ultrasonic imaging as a means of monitoring gonadal development in lumpfish (*Cyclopterus lumpus*). Physiol. Rep. 11, e15811.

Naeve, I., Mommens, M., Arukwe, A., Kjørsvik, E., 2018. Ultrasound as a noninvasive tool for monitoring reproductive physiology in female Atlantic salmon (Salmo salar). Physiol. Rep. 6, e13640.

Novelo, N.D., Tiersch, T.R., 2012. A review of the use of ultrasonography in fish reproduction. N. Am. J. Aquacult. 74, 169–181.

O’dor, R.K., Wells, M.J., 1978. Reproduction versus somatic growth: hormonal control in *Octopus vulgaris*. J. Exp. Biol. 77, 15–31.

Otero, J., González, A.F., Sieiro, M.P., Guerra, A., 2007. Reproductive cycle and energy allocation of *Octopus vulgaris* in Galician waters, NE Atlantic. Fish. Res. 85, 122–129.

Polese, G., Bertapelle, C., Di Cosmo, A., 2015. Role of olfaction in *Octopus vulgaris* reproduction. Gen. Comp. Endocrinol. 210, 55–62.

Ponte, G., Sykes, A.V., Cooke, G.M., Almansa, E., Andrews, P.R.L., 2017. The digestive tract of cephalopods: toward non-invasive *in vivo* monitoring of its physiology. Front. Physiol. 8, 403.

Reis, D., Shcherbakova, A., Riera, R., Martín, V.M., Domingues, P., Andrade, J.P., Jiménez-Prada, P., Rodríguez, C., Sykes, A.V., Almansa, E., 2021. Effects of feeding with different live preys on the lipid composition, growth and survival of *Octopus vulgaris* paralarvae. Aquac. Res. 52, 105–116.

Saber S., Macías D., Ortiz de Urbina J., Kjesbu, O.S., 2015. Stereological comparison of oocyte recruitment and batch fecundity estimates from paraffin and resin sections using spawning albacore (*Thunnus alalunga*) ovaries as a case study. J. Sea Res. 95, 226–238. 10.1016/j.seares.2014.05.003

Sánchez, F.J., Cerezo, J., García, B., 2014. Octopus vulgaris: ongrowing, in: Iglesias, J., Fuentes, L., Villanueva, R. (Eds.), Cephalopod Culture. Springer, New York, pp. 451–466.

Scherle, W., 1970. A simple method for volumetry of organs in quantitative stereology. Mikroskopie 26, 57–60.

Sieiro, P., Otero, J., Guerra, A., 2014. Contrasting macroscopic maturity staging with histological characteristics of the gonads in female *Octopus vulgaris*. Hydrobiologia 730, 113–125.

Sieiro, P., Otero, J., Aubourg, S.P., 2020. Biochemical composition and energy strategy along the reproductive cycle of female Octopus vulgaris in Galician waters (NW Spain). Frontiers in Physiology, 11, 760.

Tirado, C., Rodríguez de la Rúa, A., Bruzón, M.A., López, J.I., Márquez, L., 2003. La reproducción del pulpo (*Octopus vulgaris*) y el choco (*Sepia officinalis*) en la costa andaluza. Consejería de Agricultura y Pesca – Junta de Andalucía. Sevilla.

Wells, M.J., 1978. Octopus. Physiology and Behaviour of an Advanced Invertebrate. Chapman and Hall, New York.

Zou, W., Hong, J., Ma, Y., Yu, W., Liu, Y., Ai, C., Huang, M., Luo, X., You, W., Ke, C., 2024. Application of ultrasonography in abalone gonadal evaluation. Aquac. Fish. In press.

